# Equi-Under-Dispersed Possion Distribution (EUPoisson)

**DOI:** 10.1101/2024.10.22.619660

**Authors:** Mohamed Y. Hassan

**Affiliations:** Department of Statistics, United Arab Emirates University, P.O.Box 15551, Al-Ain, UAE

**Keywords:** Generalized Poisson, Unobserved Heterogeneity, Underdispersion, Mixtures of Weighted Distributions, Count Data

## Abstract

In this study, we propose an Equi-Under-dispersed Poisson distribution derived from a finite mixture of weighted Poisson distributions. This probability distribution is a simple, parsimonious, and flexible option suitable for modeling under-dispersed count data. It aims to overcome some of the weaknesses of existing methods in modeling Equi-Under-dispersed count data. Explicit expressions for the moment-generating function, mean, variance, and index of dispersion are derived. Real count data are used to compare its performance with that of the zero-inflated Poisson distribution and the finite mixture of Poisson distributions. Maximum likelihood estimation is implemented to estimate the parameters of the distribution, and goodness-of-fit statistical techniques are used to compare the fit of the competing distributions.

## 1 Introduction

Count data have become widely available in various scientific disciplines, including health, business, and engineering. The Poisson distribution, which is the most important discrete probability distribution, is mainly used to model and analyze datasets arising from these fields. It is employed by linking its mean to relevant covariates using the canonical (log) link function; however, some of these datasets contain unobserved heterogeneity caused by the covariates. This unobserved heterogeneity often induces overdispersion or underdispersion, and Poisson-based models cannot provide natural representations, resulting in misspecification. One of the major limitations of the Poisson distribution is that it assumes equi-dispersion (i.e., the mean and variance are equal), whereas real data from the above sources often exhibit overdispersion or underdispersion.

In recent decades, there has been extensive work to search for alternative probability distributions that could be used to model under-dispersed and over-dispersed datasets. Many authors have proposed different versions of Poisson distribution generalizations to accommodate underdispersion and overdispersion. Among them are the gamma probability model, the generalized Poisson model, and the Conway-Maxwell-Poisson (COM-Poisson) model. Some of the proposed models to handle overdispersion include negative binomial and Poisson mixtures using the EM algorithm [1,2]. [3] investigated the zero-inflated Poisson (ZIP) distribution to address the zero inflation that causes overdispersion in the count variable of the dependent variable. [4] employed the weighted Poisson distribution (WPD) as described in [5,6]; see also [7,8,9,10] for count data models. Most of the proposed models have failed to capture issues involving underdispersion. Although it has convergence problems, many researchers [11,12,13] have used the Conway-Maxwell-Poisson (CMP) distribution to fit both overdispersion and underdispersion patterns in count data. [14] discussed the generalized Poisson (GP) distribution, which can also accommodate both overdispersed and underdispersed count data. The properties of the GP distribution are discussed by [15,16,17,18]. Most recently, [19] proposed a generalized Poisson distribution suitable for modeling both overdispersed and underdispersed count data.

## 2 Finite Mixtures Models

Let *X* be a random vector of size *n* with probability density (mass) functions *f*_*j*_(*x*|*φ*_*j*_), the finite mixture of these distributions is given by

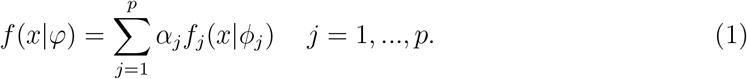

Where *f*_*j*_ is the probability density function for the jth component of the mixture, *α*_*j*_ is the mixing proportion for the jth component and *φ* is the set of the model parameters, and

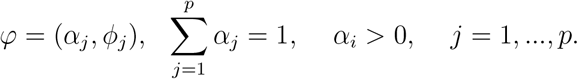

The mixing proportions are chosen by a multinomial distribution. Models derived from those distributions are flexible and allow modeling heterogeneous data. Additionally, finite mixture models have been widely applied in many areas, such as business, medicine, Vehicle crashes, and marketing.

### 2.1 Estimation

EM algorithm (Baum (1971); Dempster, Laird, and Rubin (1977)) is used for the estimation of these models. Suppose the observed vector *X* is represented by *n* observations given by (*x*_1_, …, *x*_*n*_). Let *Z* = (*Z*_1_, …, *Z*_*n*_) be unobservable random variables, where *Z*_*i*_ = (*z*_*i*1_, …, *z*_*ip*_) is a *p*-dimensional indicator vector and *z*_*ij*_ is unity if *x*_*i*_ comes from component *j* and zero otherwise. Now given all the data 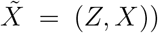), and *φ* and assuming that Z’s and *X* are independent and the *Z*_*i*_ are independent of each other, the (conditional) log-likelihood function of the whole data can be written as follows:

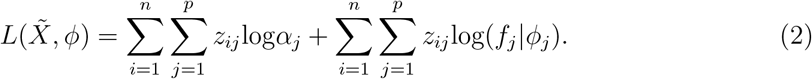

M step: Suppose that the missing *Z*_*i*_’s are now known. The estimates of the parameters *φ*_*j*_ and *α*_*j*_ can be obtained by maximizing the log-likelihood function L. The estimated parameters 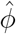 and 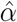, must satisfy the following M step equations

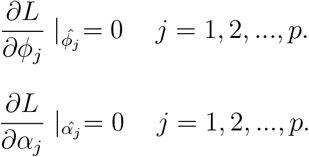

Let the conditional expectation of the *jth* component of *Z*_*i*_ be 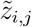 Then

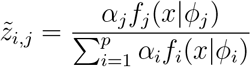

Consequently, the estimates of the model parameters can be obtained by maximizing the log-likelihood function in (2), in particular, the estimates of the mixing proportions are as follows:

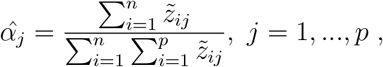

these estimates are obtained by iterating the two steps of the EM-algorithm until convergence.

## 3 Finite Mixtures of Weighted Probability Distributions

Let X be a random variable with a probability density (mass) function {*ψ*(*x, φ*), *φ* ∈ Ω} with respect to some *σ*-finite measure. Let 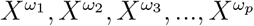be weighted versions of X with weight functions *ω*_1_(*x*), *ω*_2_(*x*), …, *ω*_*p*_(*x*) and probability density (mass) functions 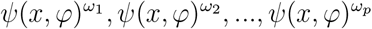 respectively. The density (mass) functions of the weighted distributions can be obtained as

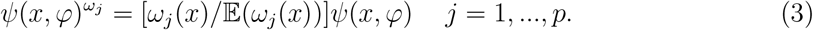

Where 𝔼(*ω*_*j*_(*x*))^′**2032**;^*s* are the expected values of the weight functions. All the weights are assumed to be positive 0 *< ω*_*j*_(*x*) *<* ∞. Consequently, the finite mixture of the weighted probability density (mass) functions 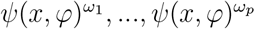 is given by

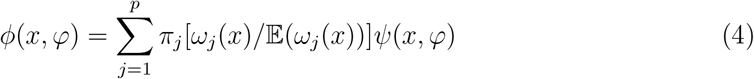

where *π, j* = 1, …, *p* are the mixing parameters with 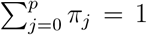 and *ψ*(*x, φ*) is the baseline density (mass) function.

### 3.1 Deriving Closed Form Probability Distributions from the Mixtures of the Weighted Distributions

If the mixing proportions 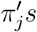 of the mixture in (4) are reparameterized as functions of the weight expectation proportional method, a unified probability density function in a closed form is obtained. There are possibly many ways to choose the reparameterizations of the mixing parameters, but the weight expectation proportional method setup allows components with larger weight expectations to have more effect on the resulted density function whereas those with lesser weight expectations have minimal impact on the distribution. If the 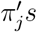 are reparameterized as follows:

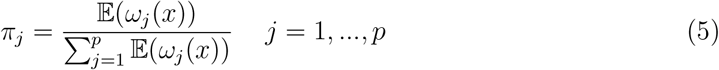

then the mixture in (4) can be rewritten as

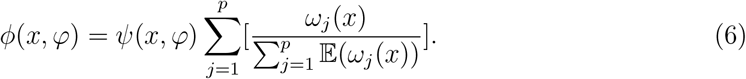

### 3.2 Distributions Derived from the Mixtures of the Weighted Poisson Distributions

If 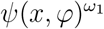 and 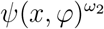 are two weighted Poisson mass functions with weights *δ* and *x*^*τ*^, where *τ* is a natural number, then by using equations (4) and (6), we obtain the following closed form probability mass function produced by the mixture of the two weighted Poisson distributions [19].

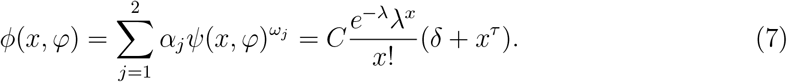

Where *C* is the normalizing constant. Other distributions derived from mixtures of weighted distributions include the probability distribution proposed by [20], which has many applications in statistics, and the Bimodal Skew-Symmetric Normal Distribution (BSSN) proposed by [21]. Since the study of underdispersed and overdispersed count data has received more attention in recent years, and most of the currently used models restrict their applications to a limited number of cases that do not explicitly consider these properties, it is necessary to investigate alternative models that could be employed for the analysis of this data. The family of distributions mentioned above explicitly takes into account the variability and shape of these datasets. A major advantage of this family is that it generates realistic dispersion indices for both underdispersed and overdispersed count data.

## 4 EquiDispersed and UnderDispersed (EUPoisson)

Definition 4.1. A random variable *X* has *EU Poisson*(*λ, δ*) distribution with parameters *λ* and *δ* if its probability mass function is given by

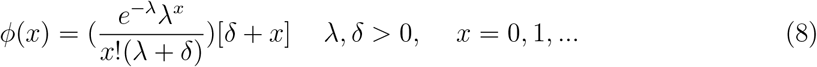

Proposition 1: If X ∼ *EUPoisson*(*λ, δ*) and if *δ* → ∞ then

(i) (*δ* + *x*)*/*(*λ* + *δ*) → 1 and
(ii) *φ*(*x, λ, δ*) → *φ*(*x, λ*) where *φ*(*x, λ*) is the Poisson distribution.

Proof:

(i) 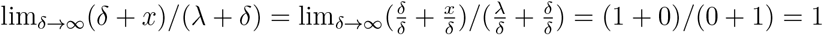 The proof of (ii) follows (i). So,
(ii) 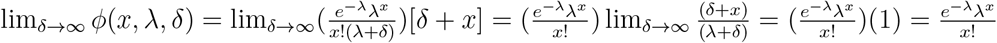

Clearly the distribution in (8) converges to the Poisson distribution as *δ* → ∞.

### 4.1 Moments and Fisher’s Index of Dispersion

The moment generating function (MGF) of the probability distribution in (8) is given by

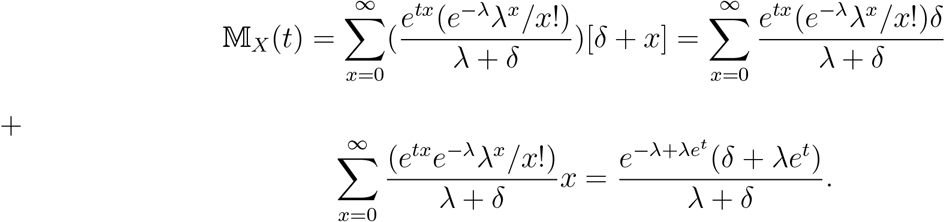

Therefore, the first and the second derivatives of the MGF are

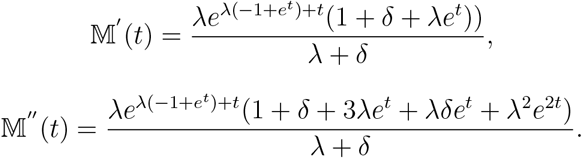

It follows that, the mean and the variance are given by

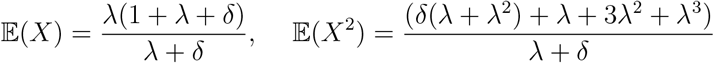

and

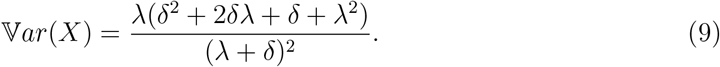

Consequently, the Fisher’s index of dispersion is given by

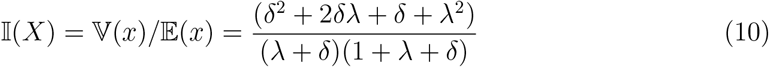

Proposition 2: If X ∼ *EUPoisson*(*λ, δ*) and

- If *δ* → ∞ then I(*X*) → 1
- If *δ* → 0 then I(*X*) → *λ/*(1 + *λ*)
- If *δ* → 0 and *λ* → 0 then I(*X*) → 0

Proof:

(i) 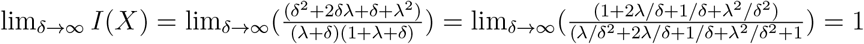
(ii) 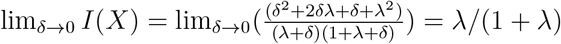
(iii) The proof of (iii) is trivial, and it follows (i) and (ii).

Figure 1 displays a three dimensional plot of the dispersion index. It can be seen that the index approaches zero at the origin and then rises until reaches its maximum, which is clearly 1.

**Figure 1:**
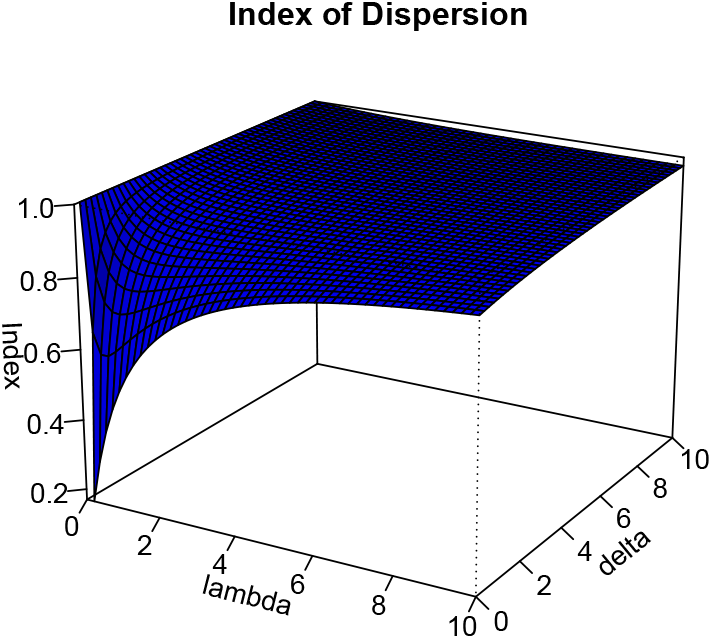
Three dimensional surface plot of the dispersion index

Figures 2, and 3 presents the plots of the dispersion index, *I*(*X*), with different values of *δ* and *λ*. The plots in the Figues show that the dispersion index converges to 1 if *δ* → ∞ or *λ* → ∞, which means that the mean and the variance are equal, see proposition 1, part (ii). For other values of *δ* and *λ*, it fluctuates in the interval [0,1). If *δ* → 0, the index is a decreasing rational function of *λ*, which is *λ/*(1 + *λ*. This function reaches its maximum at *λ* → ∞ as shown on the lower right plots of Figures 2 and 3. Other plots on the two Figues show the behaviour of the index for the given values of *λ* and *δ*.

**Figure 2:**
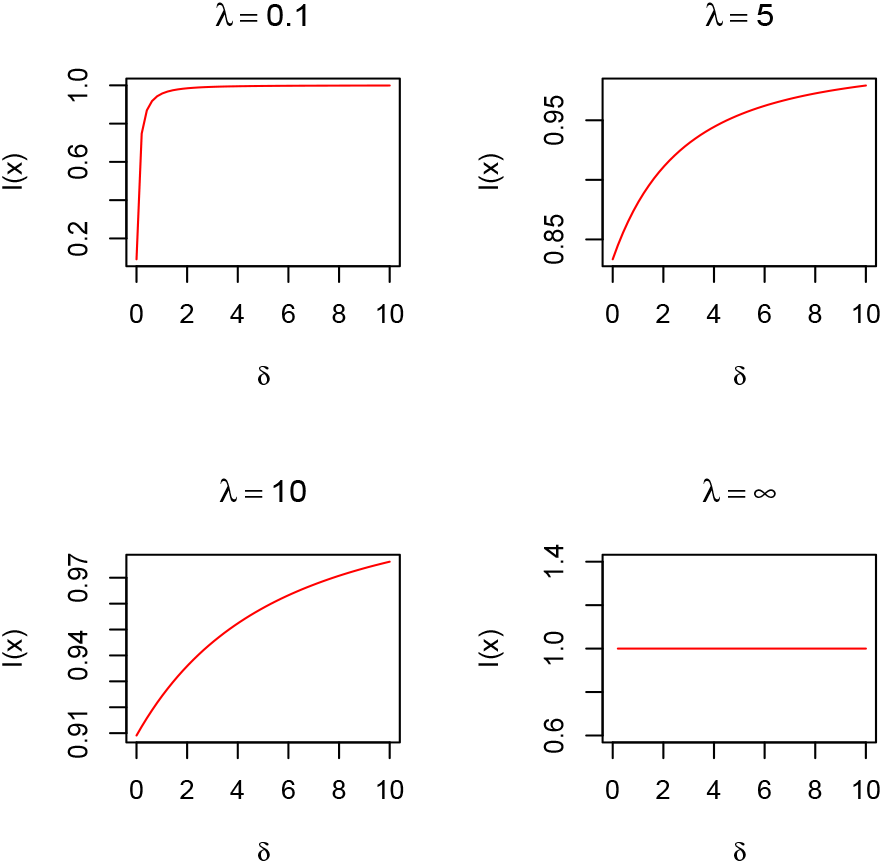
Plots of *I*(*X*) based on different values of *λ*.

**Figure 3:**
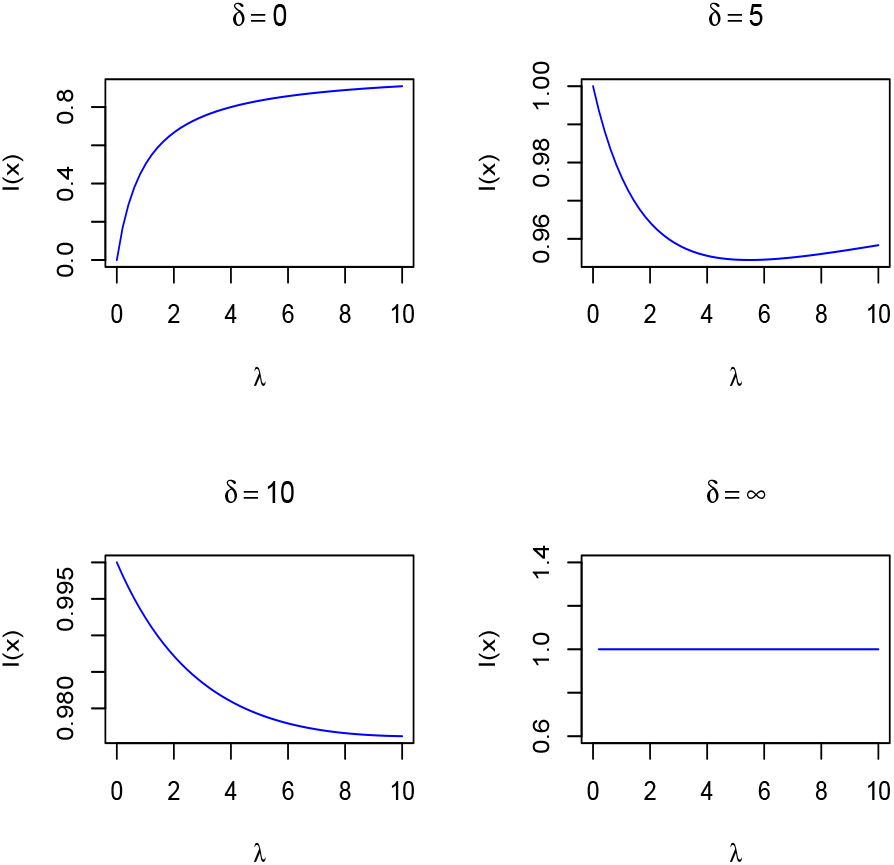
Plots of *I*(*X*) based on different values of *δ*.

## 5 Model fitting and examples

Using real data, an example is worked out in this section to illustrate the fitting of EUPoisson model and its comparison with the zero-inflated Poisson distribution and the Poisson mixture. The estimation of the parameters of the EUPoisson was carried out by maximizing the likelihood function over the parameter space *λ >* 0, *δ* ≥ 0. To avoid local maxima, the optimization routine was run with several different starting values that are widely scattered in the parameter space. Chisquare test is used for the model fitting, while the Akaike information criteria (AIC) and the Bayesian information criterion (BIC) are used for the model comparisons.

### 5.1 Cardiology Dataset

These datasets were downloaded from the University of California, Irvine (UCI) Machine Learning Repository, and they were obtained from Horton General Hospital. This data contains the number of monthly cardio-respiratory arrest transfers from the Emergency Room to Critical Care at Horton General Hospital and consists of 139 observations. The mean, variance, and index of dispersion for the dataset are 0.863, 0.728, and 0.844, respectively. From these measures, it is readily seen that the data are underdispersed. The dataset is fitted to three models: EuPoisson, Poisson mixture, and zero-inflated Poisson distributions. The bar charts of the actual data and the fitted values of the three competing models are presented in Figures 4 and 5. These figures display qualitative assessments of the fitted models on the bar charts, demonstrating the superiority of EuPoisson compared to the other two models. The EuPoisson model outperformed the other two models in three classes; it captured the second, third, and fourth classes better but did not perform well in the first class. The maximum likelihood estimates and their parameters, along with their standard errors, are given in Table 1.

**Table 1:** Model Parameter Estimates.

| Estimated Model | $\hat{\delta}$ | $\hat{\lambda}$ | $\hat{\pi}$ |
| --- | --- | --- | --- |
| **EU-Poisson** | 0.921(0.663) | 0.508(0.124) | - |
| **PoissonMix.** | - | 0.863(0.079) | 1 |
| **Zero-inflated** | - | 0.863(0.014) | 1 |

**Figure 4:**
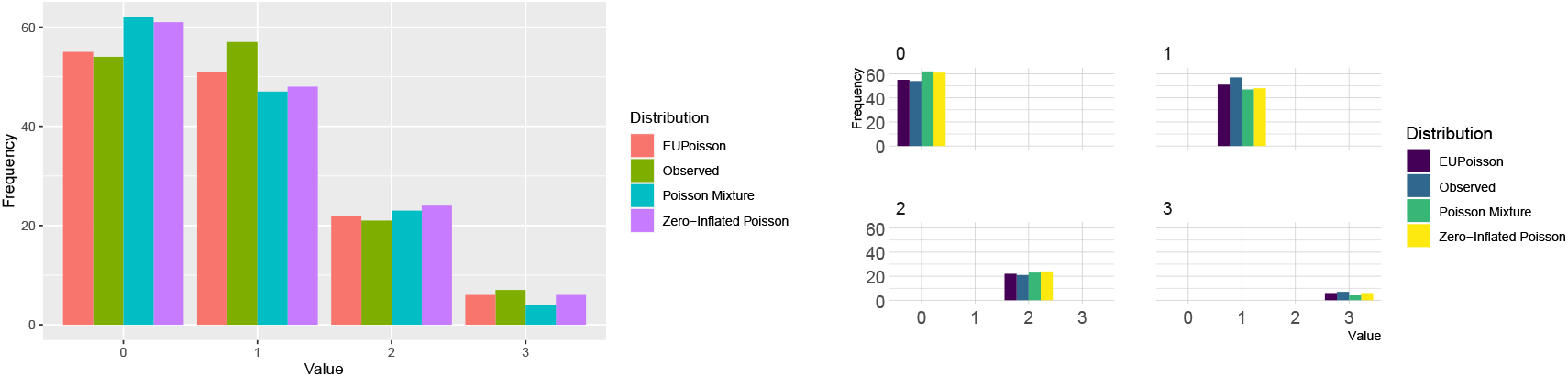
Barchart Plot of the fitted models. Figure 5: Barchart Plots of the fitted models.

The chi-square goodness-of-fit test statistic, along with its p-values and the values of the information criteria AIC and BIC, are presented in Table 2. The chi-square goodness-of-fit test statistic assesses how well the observed data fit the expected model, providing a p-value that indicates the statistical significance of the model’s fit. Lower p-values suggest that the model does not fit the data well. In addition, the Akaike Information Criterion (AIC) and the Bayesian Information Criterion (BIC) are both used to compare the relative quality of statistical models for a given dataset. AIC focuses on the trade-off between model complexity and goodness of fit, where lower AIC values indicate a better model. Similarly, BIC also penalizes model complexity but does so more strongly, making it particularly useful when the sample size is large. Together, these metrics enable a comprehensive assessment of model performance, allowing for informed decisions when selecting the most appropriate model.

**Table 2:** Model Accuracy Measures.

| **Estimated Model** | $\chi^2$ | P-value | -2LogL | BIC | AIC |
| --- | --- | --- | --- | --- | --- |
| **EUPoisson** | 0.840 | 0.657 | 327.52 | 332.47 | 328.52 |
| **PoissonMix.** | 1.285 | 0.257 | 329.47 | 339.34 | 331.47 |
| **Zero-inflated** | 1.285 | 0.257 | 329.47 | 334.42 | 330.47 |

The values presented in Table 2 summarize the performance of the three competing models. Both the zero-inflated Poisson and the EUPoisson models have two independent parameters, while the Poisson mixture model has three independent parameters. The results indicate that all three models exhibit flexibility and parsimony. However, based on the accuracy measures, the EUPoisson model demonstrates superior performance compared to the other two models.

## 6 Conclusions

The general form of the EUPoisson distribution mass function is presented and investigated. It has been shown that this distribution is capable of capturing the under-dispersed features of count data. The convergence in probability of the distribution to the Poisson distribution is demonstrated. The moment-generating function, mean, variance, and index of dispersion are derived. Some properties of the index are explored. The performance of the distribution is compared with that of the Poisson mixture and the zero-inflated Poisson using real data. The maximum likelihood method is implemented to estimate the parameters of the EUPoisson using the EM algorithm. Parameter estimates of the three models, along with their errors, are given in Table 1. The chi-square goodness-of-fit test statistics, along with their p-values and the values of the information criteria, AIC and BIC, are presented in Table 2. The results of the performance evaluations for the three models in the real data example indicate the simplicity and potential superiority of the parsimonious EUPoisson model compared with the other two well-known distributions. The theoretical properties of the index of dispersion for the EUPoisson and the outcomes of the information criteria (AIC and BIC) from the real data show that EUPoisson is the best-suited discrete distribution for the analysis of count data, and it addresses some of the weaknesses of existing models in modeling underdispersion.

